# Integrated genetic screening reveals FEN1 as a driver of stemness and temozolomide resistance in glioblastoma

**DOI:** 10.1101/2025.06.18.660339

**Authors:** Benjamin A. Brakel, Dillon McKenna, Anish Puri, Vaseem Muhammad Shaikh, Manoj Singh, Ali Saleh, Abdo-jose Tomajian, Nicholas Mikolajewicz, Marcello Beltrami, Alisha Anand, Petar Miletic, Kevin R. Brown, David Tieu, William Maich, Sabra Salim, Yujin Suk, Minomi Subapanditha, Deena M.A. Gendoo, Chitra Venugopal, Jason Moffat, Sachin Katyal, Chirayu R. Chokshi, Sheila K. Singh

## Abstract

Glioblastoma (GBM) is a lethal brain tumor with limited response to standard of care chemoradiotherapy. In this study, we conducted genome-wide CRISPR knockout screening in patient-derived glioblastoma stem cells (GSCs) to identify genetic dependencies of cell survival and therapy resistance. Our screening identified flap endonuclease 1 (FEN1) as a key driver of GSC survival, with enhanced dependency under temozolomide (TMZ) treatment. Genetic perturbation of FEN1 reduced GSC self-renewal and proliferation in vitro, and prolonged survival in a patient-derived xenograft model of GBM. FEN1 inhibition (FEN1i) preferentially affected highly aggressive or recurrent GBM models compared with less aggressive GBMs and healthy neural stem cells. Moreover, FEN1 inhibition synergized with TMZ only in these aggressive FEN1i-sensitive GSCs, providing cancer-selective killing and TMZ sensitization in the most untreatable of GBMs. Mechanistically, FEN1i-sensitive GSCs exhibited greater proliferation and sphere formation, while stalling their proliferation conferred resistance to FEN1 inhibition. Single-cell transcriptomics further linked FEN1 expression to stemness and the DNA damage response, elucidating broader determinants of FEN1 dependency. These findings establish FEN1 as a promising therapeutic target in GBM, offering a strategy for both selective targeting and enhancement of TMZ efficacy in aggressive cancers.

**Statement of Significance:** This study identifies FEN1 as a key vulnerability of glioblastoma stem cells, revealing its role in therapy resistance and stemness, and proposes FEN1 inhibition as a strategy to enhance temozolomide efficacy.

## Introduction

Glioblastoma (GBM) is the most common and aggressive primary brain tumor in adults, marked by rapid progression, therapy resistance, and dismal outcomes. Despite multimodal treatment including surgical resection, radiation therapy (RT), and temozolomide (TMZ) chemotherapy, the median survival remains less than 15 months with inevitable tumor relapse.^1^ Therapy resistance is largely attributable to the presence of glioblastoma stem cells (GSCs), a treatment-resistant and highly tumorigenic subpopulation that drives tumor growth, resistance to genotoxic therapies and recurrence. GSCs exhibit enhanced DNA repair capacity, stem-like properties, and dynamic adaptability, rendering them inherently resistant to standard of care (SoC) treatments.^2–5^

While genomic and transcriptomic profiling have offered insights into the complex molecular landscape of GBM, these static approaches fail to capture the functional drivers of GSC survival and therapy resistance. The advent of CRISPR-Cas9 technology has revolutionized functional genomics, enabling genome-wide screens to uncover essential genes in an unbiased manner.^6^ However, few efforts have focused on understanding the functional drivers of therapy resistance in GBM, particularly in the context of SoC chemoradiotherapy, leaving a gap in understanding how genetic dependencies evolve in response to standard treatments. In this study, we conducted genome-wide CRISPR knockout screens in patient-derived GSCs to systematically identify genetic vulnerabilities of GSC survival and resistance to therapy. By integrating functional genomic data across treatment conditions, we explored conditional genetic interactions of essential genes to ultimately identify dependencies that both sustain GSC viability and mediate resistance to SoC therapies. These efforts identified flap endonuclease 1 (FEN1) as a key driver of GSC survival, with heightened dependency under TMZ treatment. Further analysis revealed that FEN1 inhibition selectively targets highly aggressive, stem-like GBM cells, revealing a therapeutic vulnerability that not only disrupts GBM growth but also enhances TMZ efficacy in treatment-resistant tumors.

These findings help illuminate the genetic underpinnings of GSC survival and therapy resistance, offering a new framework for targeting functional vulnerabilities in GBM. By identifying FEN1 as a cancer-selective and therapy-sensitizing target, this work provides a basis for developing precision therapies aimed at overcoming resistance and improving outcomes for patients with this devastating disease.

## Methods

Our research complies with ethical regulations according to the Hamilton Health Sciences and McMaster Health Sciences Research Ethics Boards.

### Derivation and culture of patient-derived glioblastoma stem cell models

Primary glioblastoma (GBM) specimens and whole fetal brain samples were collected from consenting patients and families, following approval from the Hamilton Health Sciences and McMaster Health Sciences Research Ethics Board. After an initial PBS wash, specimens underwent mechanical dissociation, followed by enzymatic dissociation in PBS containing 0.013 mg/mL Liberase Thermolysin Research Grade (Millipore Sigma, #5401020001) at 37 °C for 15 minutes. The dissociated cells were filtered through a 70 µm cell strainer (Millipore Sigma, #CLS431751-50EA), centrifuged, and subjected to red blood cell lysis using a 0.8% ammonium chloride solution (STEMCELL Technologies, #07850).

Tumor cells and fetal human neural stem cells (NSCs) were cultured in the NeuroCult NS-A proliferation kit (Human) (STEMCELL Technologies, #05751), supplemented with 20 ng/mL EGF (STEMCELL Technologies, #78006), 10 ng/mL FGF (STEMCELL Technologies, #78003.2), 0.002% Heparin (w/v) (STEMCELL Technologies, #07980), and 1X antibiotic/antimycotic solution (Wisent Bio Products, #450-115-EL). During the initial two weeks of culture, cells were treated with 1X MycoZap Prophylactic (Lonza, #VZA-2031) to prevent potential mycoplasma contamination.

Cells were maintained on tissue culture-treated dishes, cell-repellent dishes (Greiner Bio, #628979), or tissue culture-treated dishes coated with Poly-L-ornithine (Millipore Sigma, #P4957) and mouse Laminin (Corning, #354232). All cultures were kept at 37 °C in a 5% CO environment. Regular mycoplasma testing was performed, and if necessary, cells were treated with an additional two-week course of 1X MycoZap.

### Genome-wide CRISPR-Cas9 knockout screening

For our in-house screens, the Toronto KnockOut version 3.0 (TKOv3) guide RNA (gRNA) library (Addgene no. 90294) was packaged into lentivirus and multiplicity of infection (MOI) was determined as previously reported.^7^ Our dropout screen (BT935) was performed with methodology as described by our group previously.^8^ gRNA read counts were acquired from additional in-house screens (BT972, BT241, and BT594)^8^ and others (G440_L, G523_L, G729_L, G532_L, G620_L, BT67_L, G361_L, G691_L, G809_L, G549_L, G583_L, CB660, U5, HF7450, HF6562).^9–11^ For our drug-treated screen (BT935), infected and puromycin-selected cells were split into one of four treatment arms: control dimethyl sulfoxide (DMSO) treatment, sublethal IC_20-30_ radiotherapy (RT), sublethal IC_20-30_ temozolomide (TMZ), or sublethal IC_20-30_ RT and TMZ. For the combined RT and TMZ arm, each plate was treated with TMZ (Millipore Sigma no. T2577) one hour prior to RT (Faxitron RX-650). For the entire duration of the screen, treatments were administered in cycles of 5 days on and 2 days off to allow for recovery and passaging. Library preparation, Illumina sequencing of gRNAs, mapping and quantification of gRNAs, quality control analysis and gRNA scoring were performed as previously described.^7^ In summary, to quantify gene fitness in each screen, log fold change (LFC) of each gRNA was quantified by comparing read-depth-normalized gRNA counts at the beginning (T0) and end (Tn) of each screen. LFC for all gRNAs targeting a single gene were averaged, with four gRNAs targeting each gene in TKOv3.^7^ Core essential gene sets (reference)^6,7,12^ and non-essential gene sets (background)^6,7^ were used to asses quality control.

For dropout screens, LFC values of core essential genes and non-essential genes were used to apply the Bayesian Analysis of Gene Essentiality (BAGEL) algorithm^13^ to determine gene fitness. A Bayes Factor (BF) score of ≥ 5 and false discovery rate (FDR) < 0.05 were used to identify genes essential for cell survival. GBM-enriched genetic dependencies were identified as genes essential across all GBM models, after excluding genes from core essential gene datasets and those essential in a majority of NSC lines.^7,12^ For the drug-treated screen (BT935), LFC of normalized gRNA read counts were compared between control (DMSO) and treatment (RT + TMZ) arms.

### Knockout and knockdown generation

CRISPR knockout gRNA targeting AAVS1 (5’-GGGGCCACTAGGGACAGGAT-3’), FEN1 (5’-TCTGAGGAGCGAATCCGCAG-3’) or TP53 (5’-GCAGTCACAGCACATGACGG-3’) were obtained from TKOv3^7^ and cloned into the single-gRNA lentiCRISPRv2 construct (Addgene, 52961). For knockdowns, hairpin oligonucleotides were constructed targeting the coding sequence of GFP (5’-TGACCCTGAAGTTCATCTGCA-3’) or FEN1 (5’-GCAGTGACTACTGTGAGAGTA-3’) and cloned into the pLKO.1 construct (Addgene, 10878). Constructs were validated using Sanger sequencing. Each construct was packaged into lentivirus using second-generation packaging vectors (psPAX2 and pMD2.G with PEI transfection reagent) and tumor cells were infected as described above at an MOI of 1. Twenty-four hours after infection, virus-containing media were replaced with fresh medium containing puromycin (1–3 µg/ml), and cells were incubated for an additional 24–72 h.

### In vitro assays for self-renewal and proliferation

#### Cell preparation

Tumor cells or NSCs, with or without genetic knockout, were dissociated into a single cell suspension using incubation with TrypLE (ThermoFisher Scientific no. 12605028) for 5 minutes at 37 °C. After washing with PBS, the cell suspension was filtered through a 35 µm cell strainer and plated for assays. All assays were performed using NeuroCult NS-A proliferation kit (Human), supplemented as described above.

#### Secondary sphere formation assays

Cells were plated at 200 cells/well in flat-bottom tissue culture-treated 96-well microplates. Cells were cultured in 200 µL of media and incubated for 7 days at 37 °C and 5% CO_2_. Following incubation, cell spheres were counted by microscopy. An unpaired *t*-test was used to compare between samples.

#### Proliferation assays

Cells were plated at 1000 cells/well in flat-bottom tissue culture-treated 96-well microplates. Cells were cultured in 200 µL of media and incubated for 7 days at 37 °C and 5% CO_2_. Compounds, if used, were added at the desired concentrations, maintaining a constant volume of vehicle across treatments/control. All experiments were conducted with three to five experimental replicates per sample. To quantify proliferation, 20 µL of PrestoBlue Cell Viability Reagent was added to each well followed by incubation for 4 hours at 37 °C and 5% CO_2_. Fluorescence was measured on a FLUOstar Omega Fluorescence 566 Microplate reader (BMG LABTECH) (excitation 540 nm and emission 570 nm) and analyzed using Omega software. An unpaired *t*-test was used to compare between samples. For FEN1 inhibitor dose response assays, cells were treated with FEN1 inhibitor or DMSO (vehicle control) once and incubated for 7 days before readout. For TMZ dose response assays, cells were treated with TMZ or DMSO for 5 consecutive days at the same dose and assay read out was 7 days after plating. Paclitaxel (PTX) assays were done in similar fashion, after culturing cells with 40 nM PTX or control DMSO for 6 days prior to plating the assay, with readout occurring 7 days after plating the assay.

### In vivo xenograft model

Tumor cells were orthotopically injected into immune-compromised mice as previously described.^2^ In summary, NOD-SCID mice aged 6–12 weeks were anesthetized with isoflurane gas (5% for induction, 2.5% for maintenance). A total of 100,000 tumor cells from cultured cell lines were suspended in 10 µl of PBS and administered into the right frontal lobe of each mouse. Tumor presence was verified via magnetic resonance imaging. Mice were euthanized upon reaching humane endpoint or experiencing a 20% weight loss. Survival analysis was performed in R using the survfit() and surv() functions from the ‘survival’ package (version 3.2-3) and visualized with the ‘survminer’ package (version 0.4.8).

### Synergy assays

Cells were plated and treated as described for proliferation assays, with increasing TMZ and FEN1 inhibitor concentrations while keeping a constant DMSO (vehicle) concentration. Expected inhibition was calculated using bliss additivity and synergy was calculated using bliss excess as described previously^14^ using SynergyFinder (v2.0).

### Immunoblotting

For western blotting, cell pellets were thawed and lysed immediately. Protein extracts were prepared using lysis buffer (50 mM Tris-HCl, 200 mM NaCl, 0.2% NP-40, 1% Tween-20 (v/v)) supplemented with 1 mM NaF, 1 mM sodium vanadate, 50 mM βglycerophosphate, 2 mM PMSF and Complete protease inhibitor (Roche). Lysates were then centrifuged at 13,000 rpm for 10 min and supernatant was collected. Protein concentration was determined by Bradford assay (Bio-Rad), proteins (20 µg per lane) were separated through a Bolt 4–12% (w/v) Bis-Tris plus gel (Life Technologies) and transferred onto nitrocellulose membrane (Bio-Rad). Blots were sequentially immunostained with the antibodies indicated in Supplementary Table 1. For secondary antibodies: blots were subsequently incubated with either horseradish peroxidase– conjugated goat anti-mouse (Bio-Rad, cat# 1706516, RRID: AB_2921252, 1:2000) or goat anti-rabbit (Bio-Rad, cat# 1706515, RRID: AB_11125142, 1:2000) and detected using Clarity chemiluminescence reagent (BioRad) and imaged using a ImageQuant LAS 500 automated chemiluminescence imager (GE Healthcare). Densitometry analysis of individual protein band signals was performed via ImageJ.

### Comet assay

Tumor cells were dissociated into a single-cell suspension using 5 μL Liberase Blendenzyme (0.2 Wu nsch unit/mL, Sigma, CAT# 5401119001) in 1mL PBS for 5 min at 37 °C. Cells were plated at a density of 3000-5000 cells/well in 100 μL of culture media in a 96-well Ultra-Low Attachment Surface plate (Costar®, Corning, CAT#3474). After 48 hrs of treatment with Temozolomide (0-200 μM) and/or FEN1 inhibitor (30 μM), cells were collected, embedded in low-melting agarose, and transferred onto 96-well comet slides. The slides underwent lysis to remove membranes, an alkaline buffer to unwind the DNA, and then electrophoresis at 4 °C for 40 minutes at 21V, after which slides were washed, dried, and stained with Sybr Green. DNA damage was visualized using a Cytation V cell imaging reader and analyzed with Gen 5 software for comet tail moment as indicator for DNA damage.

### KI67 immunofluorescence

Tumor cells were cultured in cell culture chamber slides until they reached approximately 70–80% confluence. The cells were then fixed with 10% formalin for 20 minutes at room temperature, followed by three washes with PBS. Permeabilization was performed using 0.1% Triton X-100 in PBS for 15 minutes at room temperature. After an additional three washes with PBS, cells were blocked with 5% bovine serum albumin (BSA) in PBS for 1 hour at room temperature to prevent nonspecific binding.

Cells were incubated overnight at 4 °C with a primary antibody against Ki-67 (Rabbit anti-Ki67 antibody, Thermo Fisher Scientific, Catalogue #RM-9106-S0, RRID: AB_2341197) diluted at 1:1000 in 1% BSA/TBS. The following day, cells were washed three times with PBS and incubated with a fluorophore-conjugated secondary antibody (Goat anti-Rabbit IgG (H+L) Alexa Fluor 488, Thermo Fisher Scientific, Catalogue #A-11008, RRID: AB_143165) diluted at 1:500 in 0.1% BSA/TBS for 2 hours at room temperature in the dark. After washing three times with PBS, nuclei were counterstained with 4′,6-diamidino-2-phenylindole (DAPI) for 5 minutes. Coverslips were mounted and sealed.

Fluorescent images were acquired using a fluorescence microscope with 20X magnification. Image processing and quantification were performed using ImageJ.

### scRNA-seq analysis

scRNA-seq data from Abdelfattah et al. (2022)^15^ was obtained from Gene Expression Omnibus (GEO; accession number GSE182109) and Wang et al. (2022)^16^ from GEO (accession number GSE131928). scRNA-seq datasets were normalized using the *Seurat* (v 4.2.1) workflow.^17^ In brief, count matrices for each tumor were loaded into separate Seurat objects and normalized using NormalizeData function (…) with default parameters. Normalized values were used for gene expression and correlation analyses.

The curated list of gene sets that was used for gene program scoring included HALLMARK,^18^ Gavish hallmark cancer programs,^19^ Greenwald GBM subtypes,^20^ Neftel GBM subtypes,^5^ Verhaak GBM subtypes,^21^ CancerSEA,^22^ and IVY spatial signatures,^23^ along with select stemness-associated gene sets, including StemSig,^24^ iPSC,^25^ Nanog/Sox2/ES1,^26^ and StemCell.^27^ We also calculated additional stemness indices using Correlation of Connectome and Trascriptome (CCAT^28^) and StemSC.^29^ Gene set activities were calculated using the AddModuleScore(…) function in Seurat,^17^ which corrects for baseline expression to account for technical and biological variability.

Spearman Correlations between each gene program and FEN1 expressions were calculated within each patient tumor and averaged across all tumors. Significance for correlations and pairwise-expression comparisons was determined using Wilcoxon tests followed by post-hoc Benjamini-Hochberg corrections.

To evaluate FEN1 gene-level associations, the normalized co-dependency index (nCDI) was calculated at the tumor-level between FEN1 and each gene method using the FindCDIMarkers (…, features.x = “FEN1”) function with default parameters from the *scMiko* R package (v. 1.0.0).^30^ CDI scores across all tumors were then averaged and rank-ordered to highlight the top FEN1-associated genes. FEN1-associated genes were defined as those that satisfied the following inter- and intra-tumoral significance criteria: 1) average of all tumor-level nCDI scores was significantly greater than zero, at 5% false discovery rate, and 2) tumor-level nCDI was significant at 5% false discovery rate in at least 50% of tumor samples.

### Statistical analysis

All experimental data is accompanied by the number of experimental replicates in the figure legends or text. Unless stated otherwise, statistical significance was assessed using an unpaired Student’s *t*-test, with *p<0.05, **p<0.01, ***p<0.001, ****p<0.0001. Statistical analyses were performed using R.

### Data availability

All processed data is included in the manuscript and supplemental materials. All custom code used is available on GitHub (https://github.com/NMikolajewicz/Brakel-2025).

### Biological material availability statement

All unique biological resources are available upon request from the corresponding author.

## Results

### Genome-scale identification of genetic dependencies

To identify genetic drivers of GSC survival in a cancer-specific context, we conducted pooled, genome-wide CRISPR-Cas9 loss-of-function screening (Figure 1A, Supplementary Figure 1) in a patient-derived GSC model (BT935), and compiled data from an additional fourteen GSC models and four neural stem cell (NSC) models screened in our lab and by others.^8–11^ Beginning with genes essential across all GSC lines (Bayes factor ≥ 5, FDR < 0.05), those belonging to core essential gene sets^7,12^ and essential in a majority of NSC models were excluded, resulting in thirty three genes of interest (Figure 1B, Supplementary Table 2). The relative lack of GBM-enriched genetic dependencies highlights the large overlap of functional drivers between NSCs and GSCs, and the significant heterogeneity between GBM models. The resulting GBM-enriched genetic dependencies were involved in several tumorigenic processes such as DNA replication and repair (FEN1 and WAPAL), RNA splicing (LSM6 and WBP11), nuclear pore complex transport (NUP85), protein translation/ribosomal function (EIF3G, EEF1G, EIF3E, NELFA, RRS1, MRPL53, MRPS23), and proteasomal degradation (PSMA1). These findings are indicative of a shift towards a more proliferative state of GSCs with increased metabolic demands, elevated protein synthesis and proteotoxicity compared to NSCs.

**Figure 1.**
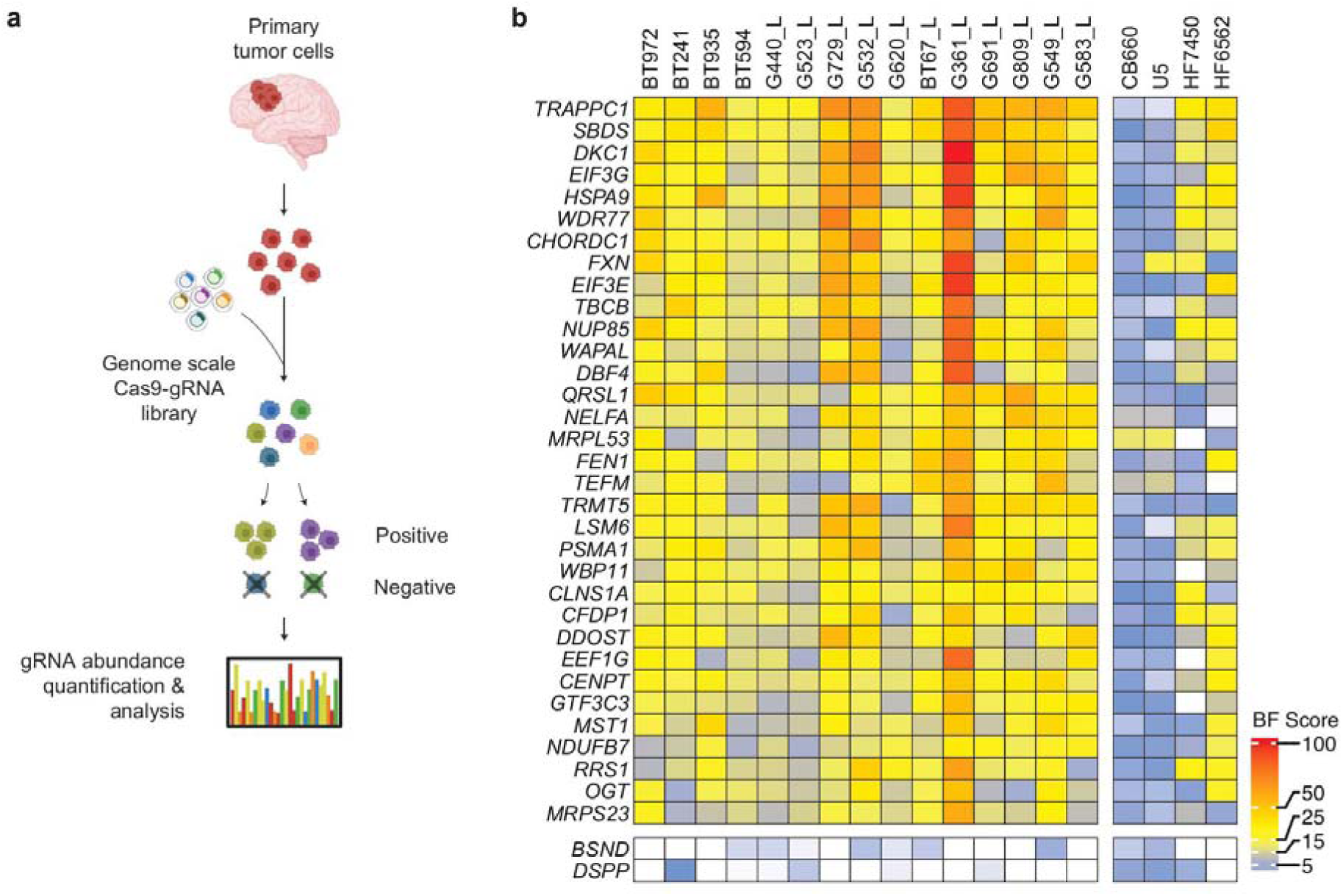
Genome-scale identification of GBM-enriched genetic dependencies. (A) Schematic of genome-scale CRISPR-Cas9 knockout screening used to identify genetic dependencies in patient-derived GBM and NSC models. Cells were infected with a genome-wide CRISPR-Cas9 gene knockout library and cultured over 15-18 doublings. Guide RNA (gRNA) fold-change within cell populations at the beginning and end of each screen were used to determine gene essentiality. (B) Log_2_ (fold change) of normalized guide RNA read counts for GBM-enriched genetic dependencies across fifteen patient-derived GBM models and four NSC models.

### FEN1 drives GSC proliferation and therapy resistance

We next investigated how these genetic dependencies contribute to therapy resistance. Essential genes elude conventional screens for therapy-sensitizing targets, as their loss impairs proliferation regardless of treatment. Consequently, their role in therapy resistance remains largely unexplored. To address this, we integrated the previous untreated screening data with chemoradiotherapy genetic screening, ranking GBM-enriched dependencies based on their differential effects between treated and untreated conditions. This approach enabled the identification of essential genes contributing to GSC therapy resistance.

Using the patient-derived GSC model BT935, we performed a genome-wide CRISPR-Cas9 loss-of-function screen under four conditions: vehicle control (DMSO), sublethal radiation (RT), sublethal temozolomide (TMZ), or combination therapy (Figure 2A). Notably, fewer than 25% of conditional genetic interactions were shared across treatment arms, underscoring the specificity of resistance mechanisms and heterogeneity among drivers of genotoxic therapy resistance.

**Figure 2.**
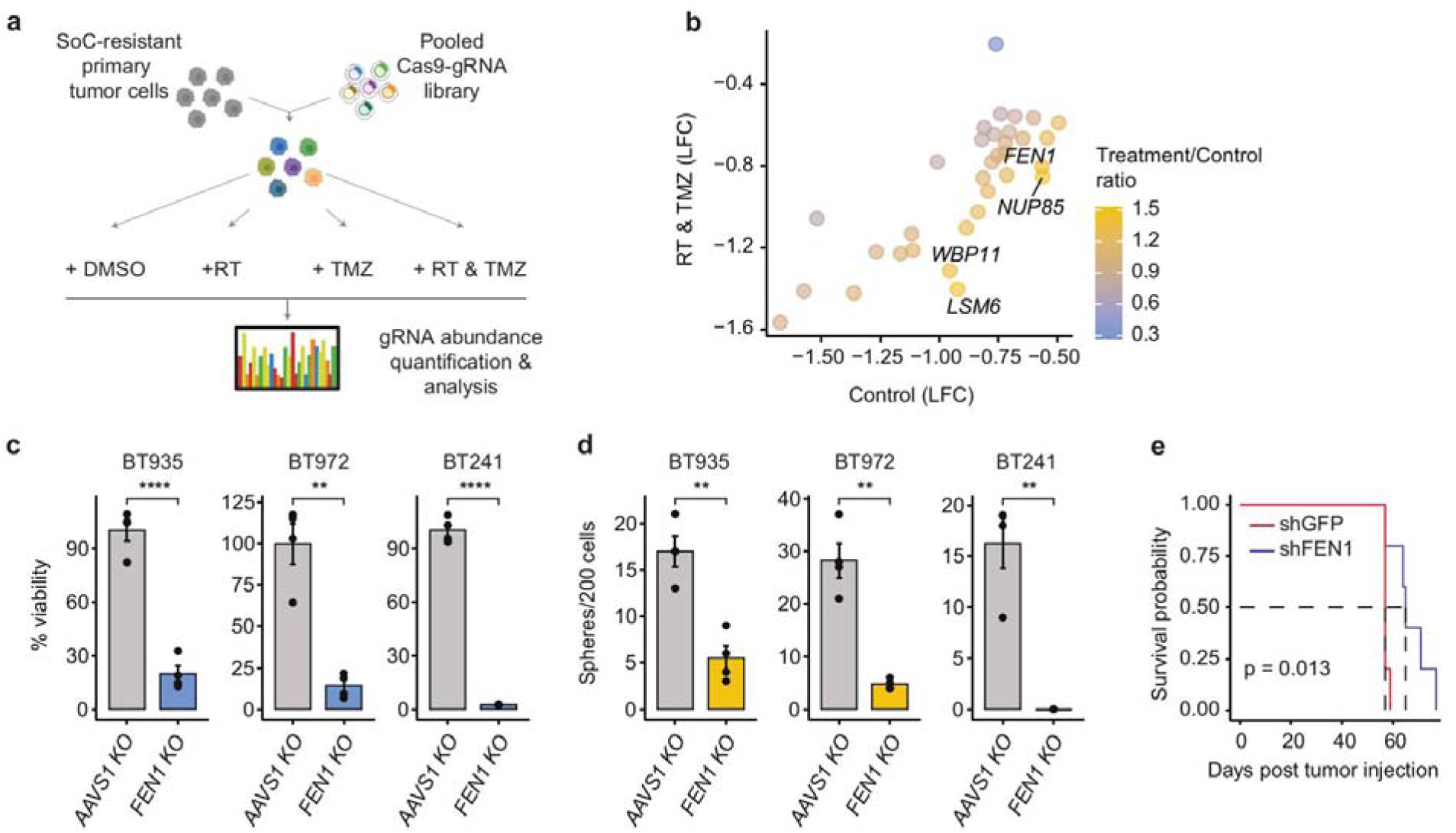
Identification of FEN1 as a driver of GSC proliferation and therapy resistance. (A) Schematic of chemoradiogenetic screen used to identify genetic dependencies of therapy resistance in GBM model BT935. Cells were infected with a genome-wide CRISPR-Cas9 gene knockout library and exposed to DMSO, sublethal RT, sublethal TMZ or combination treatment (RT & TMZ). gRNA log_2_ (fold change) (LFC) within cell populations at the beginning and end of each screen were used to determine gene essentiality. (B) Comparison of gene-level LFC between DMSO and combination treatment arms for the thirty-three previously identified GBM-enriched genetic dependencies. Node color corresponds to differential fitness ratio (LFC SoC / LFC DMSO), and genes with the highest enrichment in the therapy-treated arm are highlighted. (C, D) Evaluation of FEN1 knockout on (C) proliferation and (D) sphere formation of GSCs, compared with a gRNA targeting AAVS1 (control). Data are represented as mean ± s.e.; *n* = 4 experimental replicates; *P* values from unpaired *t*-test (\*\**P* < 0.01 and \*\*\*\**P* < 0.0001). (E) Kaplan–Meier survival analysis of immunocompromised mice engrafted with GBM tumor cells with FEN1 or GFP (control) knockdown (n = 5 per knockdown). *P* value from log-rank (Mantel–Cox) test.

To identify genes affecting fitness upon therapy exposure, we compared normalized guide RNA read counts between SoC-treated and untreated arms using differential fitness ratios (LFC SoC / LFC DMSO) (Figure 2B, Supplementary Table 3). The top candidates were relatively unexplored in GBM and included genes involved in several processes traditionally associated with resistance to genotoxic therapies such as DNA repair (MMR, HR), nuclear transport (nuclear pore complex), and RNA splicing. Analysis of TCGA data^31^ revealed significantly prolonged median survival in low grade glioma patients with low expression of several top ranked genes, including flap endonuclease 1 (FEN1) (HR (high-expressing) = 1.9, *P* = 0.01; Supplementary Figure 2A).

A key player in DNA replication and repair, FEN1 cleaves the 5’ ends of Okazaki fragments and processes DNA damage through cleaving 5’ overhanging nucleotides. It has also been implicated in apoptosis, telomere maintenance, and other important cellular functions, highlighting its role in genomic maintenance and stabilization.^32,33^ Perturbation of FEN1 has been shown to sensitize tumor cells to DNA-methylating agents across multiple cancers, posing it as a therapeutically relevant target.^34–36^ While complete knockout of FEN1 is embryonically lethal, heterozygous loss does not show severe effects, indicating a potential therapeutic window.^37^ Notably, FEN1 is highly expressed in proliferative tissues and cancers, correlating with advanced tumor grade and aggressiveness.^36,38–42^ Given its role in homologous recombination and base excision repair—critical pathways for repairing TMZ-induced DNA damage—FEN1 emerges as a promising therapeutic target for overcoming resistance.^43,44^

Supporting our genomic observations and validating the effect of FEN1 perturbation on stem-like properties of GSCs, CRISPR-Cas9 knockout of FEN1 significantly reduced proliferation and secondary sphere formation (measures of cell stemness) across several patient-derived GBM models (Figures 2C and 2D; *P* < 0.05, unpaired *t*-test, Supplementary Figure 2B). Furthermore, while complete knockout is not feasible *in vivo* due to FEN1 essentiality, genetic knockdown of FEN1 significantly extended the median survival of a patient-derived xenograft model of a highly aggressive, recurrent GBM model (Figure 2E; *P* < 0.05, Supplementary Figure 2C).

### FEN1 dependency extends to TMZ resistance

To explore the therapeutic window of FEN1 perturbation, we acquired a small molecule inhibitor of FEN1 (FEN1-in-4, Compound 2^45^). FEN1 inhibition stratified six patient-derived GBM models and two healthy NSC models into two distinct phenotypes (Figure 3A). Three GBM models showed sensitivity to FEN1 inhibition (herein called “FEN1i-sensitive”; BT972, BT241, and BT935), while three GBM models and the two NSC models showed resistance (herein called “FEN1i-resistant”; BT594, MBT357, MBT373, NSC195, NSC201). Notably, the GBM models sensitive to FEN1 inhibition appeared to be the more aggressive models in our experience, including the two recurrent GBM models assayed.

**Figure 3.**
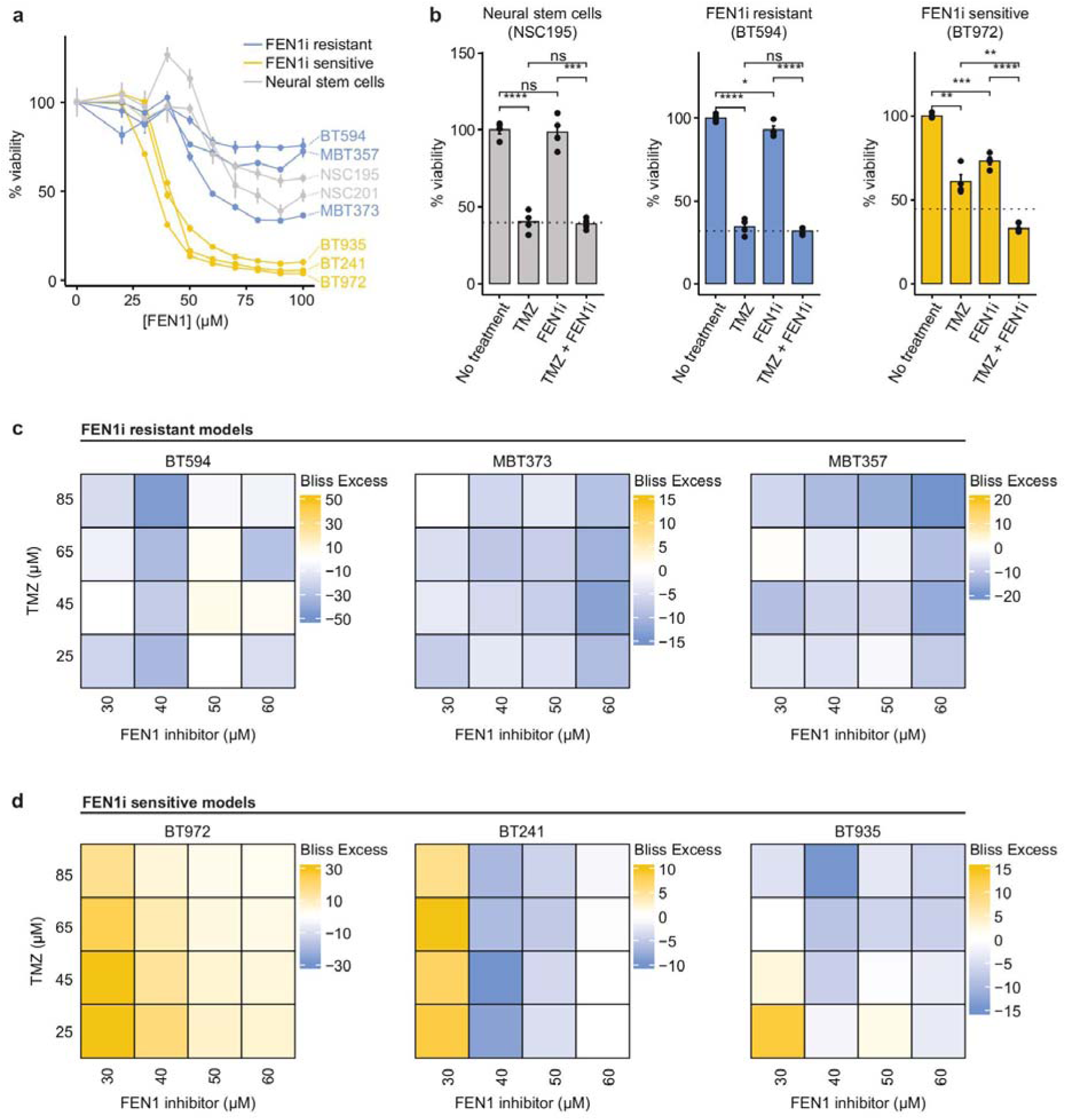
FEN1 dependency extends to TMZ resistance. (A) Cell viability assays of GSC and NSC lines exposed to increasing concentrations of FEN1 inhibitor over 7 days. Data are represented as mean ± s.e.; n = 4 experimental replicates. (B) Cell viability assays of NSC, FEN1i-resistant and FEN1i-sensitive GSC lines exposed to DMSO (control), 25 μM TMZ, 30 μM FEN1i or combination (TMZ & FEN1i). Dashed lines represent expected viability under combination treatment. Data are represented as mean ± s.e.; n = 4 experimental replicates. (C, D) Synergy assays between TMZ and FEN1 inhibitor in (C) FEN1i-resistant and (D) FEN1i-sensitive GBM models. Colors correspond to bliss excess synergy scores wherein more positive values indicate stronger synergy; n = 4 experimental replicates.

To validate whether FEN1 contributes to GSC therapy resistance, we focused on TMZ over RT due to the pertinent role of FEN1 in TMZ response pathways. Additionally, there was a greater reliance on FEN1 in the TMZ-treated arm of our screen compared to the RT-treated arm, and there was no correlation between sensitivity to FEN1 inhibition and TMZ indicating the two agents are suitable for combination with no cross-resistance (Supplementary Figures 3A and 3B). Thus, we explored the effects of FEN1 inhibition on TMZ-induced cell death in both FEN1i-sensitive and FEN1i-resistant GSCs. We began by using a patient-matched primary-recurrent pair of GBM models, where the primary GBM (BT594) was FEN1i-resistant while the more aggressive, treatment-resistant recurrent GBM (BT972) was FEN1i-sensitive. Consistent with its prior exposure to SoC chemoradiotherapy, the recurrent GSCs were highly resistant to a therapeutically relevant concentration of TMZ compared to the primary counterpart and healthy NSCs (Figure 3B).^46–48^ In contrast, this model was more sensitive to FEN1 inhibition, suggesting FEN1 dependency may have been acquired upon recurrence, and was the only model that FEN1 inhibition sensitized to TMZ (*P* < 0.05, unpaired *t*-test). This prompted us to investigate this synergy across all six GBM models previously assayed. While no synergy with TMZ was present in the three FEN1i-resistant GBM models (Figure 3C), there was strong sensitization to TMZ-induced killing at low concentrations of FEN1 inhibitor across all three FEN1i-sensitive GBM models, indicated by synergistic cell death with high bliss excess (Figure 3D). These results show that FEN1 dependency extends to therapy resistance, selectively in GSCs that are highly dependent on FEN1 prior to treatment-exposure, and positions FEN1 as a cancer-selective and therapy-sensitizing target for a subset of highly aggressive GBMs.

### FEN1 inhibition mediates TMZ sensitization through increased DNA damage

We next explored this sensitization to TMZ at the level of DNA damage and DNA damage repair. Upon exposure to high dose (100 µM) TMZ, FEN1 protein expression was significantly higher in FEN1i-sensitive GSCs (Figure 4A, Supplementary Figure 4A). Comparing the patient-matched primary-recurrent GBM models, TMZ treatment increased FEN1 protein expression only in the recurrent GSCs, suggestive of its reliance on FEN1 when exposed to TMZ (Figure 4B). To determine how FEN1 inhibition enhances TMZ activity, we proceeded to quantify DNA double stranded breaks (DSBs) and assay the expression of several DNA damage response proteins involved in DSB repair after exposure to TMZ, FEN1 inhibition or combination treatment. Using a comet assay to quantify DSBs, exposure to 30 µM FEN1 inhibitor did not induce DSBs in either GBM model (Figure 4C and 4D, Supplementary Figure 4B). However, this concentration of FEN1 inhibitor did enhance DSBs induced by increasing concentrations of TMZ, markedly more so in the recurrent GBM, confirming that inhibition of FEN1 sensitizes GSCs to TMZ through enhancing DNA damage. Looking at DNA damage repair proteins, changes in phosphorylated (active) Chk1, Chk2 and ATM mimicked our functional findings (Figure 4E and 4F, Supplementary Figure 4C and 4D). While the FEN1i-resistant GSCs showed a significant DNA repair response to TMZ and no response to FEN1 inhibition, the FEN1i-sensitive GSCs showed a greater response to FEN1 inhibition which carried over to the combination treatment arm. Interestingly, in contrast to a previously identified synthetic lethality between FEN1 and DNA protein kinase (DNA-PK) in glioma, changes in phosphorylated DNA-PK protein levels did not discriminate between the primary and recurrent GBM (Supplementary Figure 4D) indicating that the effects of FEN1 inhibition are independent of non-homologous end joining (NHEJ). Confirming these findings across multiple GBM models, Chk1 and Chk2 phosphorylation was significantly higher in FEN1i-sensitive GSCs after treatment with 100 µM TMZ compared to FEN1i-resistant GSCs (Figure 4G). These data support that FEN1 inhibition targets this subset of GBMs and enhances TMZ efficacy through enhancing TMZ-induced DSBs.

**Figure 4.**
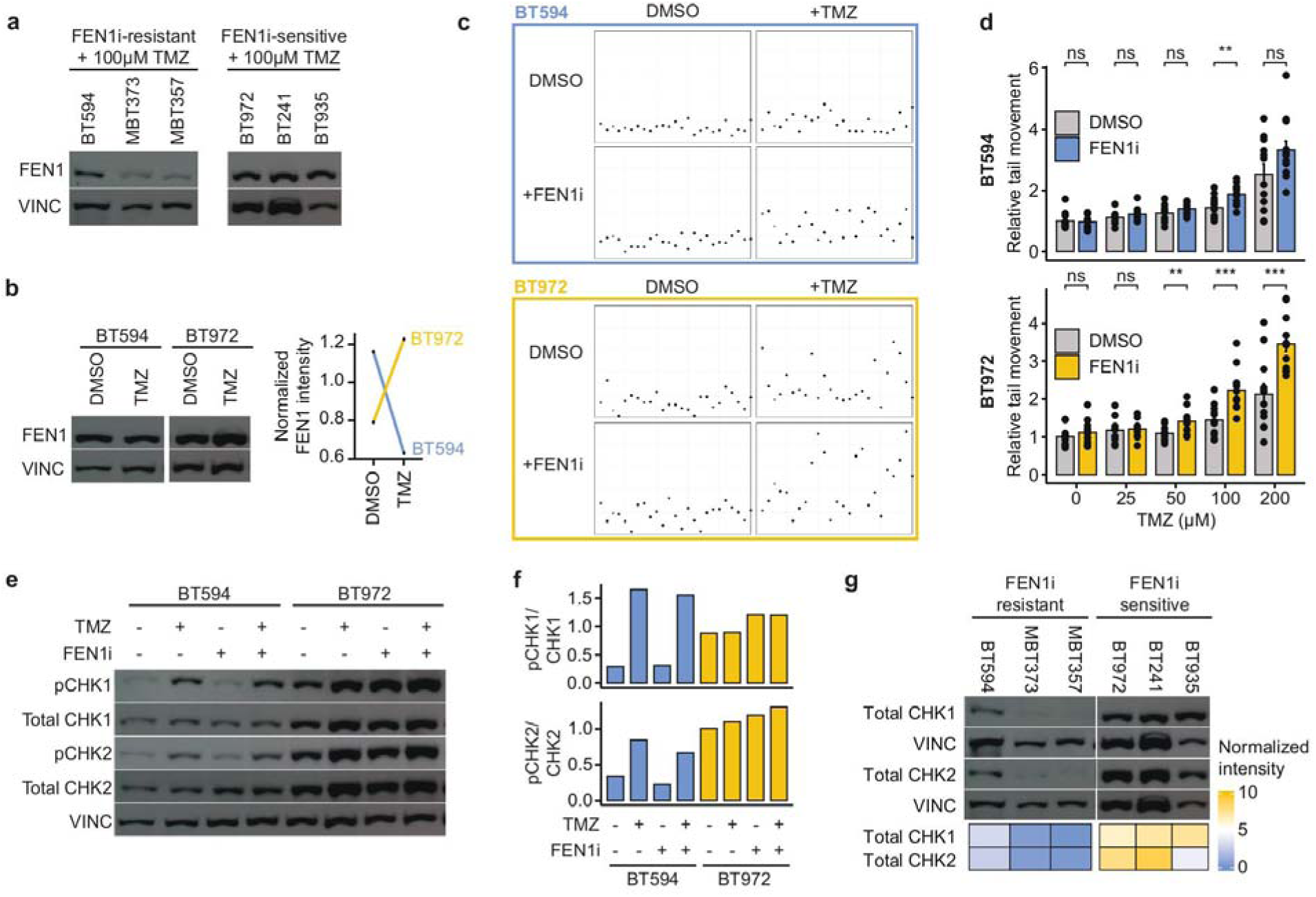
FEN1 inhibition enhances TMZ-induced DNA damage. (A) Immunoblotting of FEN1i-resistant and sensitive GSCs for FEN1 protein after treatment with 100 µM TMZ. (B) Immunoblotting of patient-matched primary (FEN1i-resistant; BT594) and recurrent GSCs (FEN1i-sensitive; BT972) for FEN1 protein after treatment with DMSO (control) or 100 µM TMZ. FEN1 signal intensity normalized to vinculin control is shown. (C) Scatterplots indicating representative individual cellular comet tail moment values of FEN1i-resistant (BT594) and sensitive (BT972) GSCs from each corresponding cell/treatment type: DMSO alone (control), TMZ (200 µM), FEN1i (30 µM) or combination (TMZ/FEN1i, 200 µM/ 30 µM). (D) Quantification of comet assays. Relative tail movement of FEN1i-resistant and sensitive GSCs is shown with increasing concentrations of TMZ ± 30 µM FEN1 inhibitor for 48 hours. Data are represented as mean ± s.e.; n = 12 experimental replicates. (E) Immunoblotting of FEN1i-resistant (BT594) and sensitive (BT972) GSCs for phosphorylated CHK1, total CHK1, phosphorylated CHK2 and total CHK2 after 3-day treatment with DMSO (control), TMZ (45 µM), FEN1i (30 µM) or combination (TMZ & FEN1i). (F) Quantification of immunoblotting in (E). Proportion of phosphorylated (active) CHK1 and CHK2 are shown, with signal intensities normalized to vinculin control. (G) Immunoblotting of FEN1i-resistant and sensitive GSCs for total CHK1 and total CHK2 protein after treatment with TMZ (100 µM). Heatmap color corresponds to normalized FEN1 intensity.

### FEN1 maintains GSC stemness and proliferation

We next sought to characterize the determinants of FEN1 dependency. Comparing stemness properties, FEN1i-resistant GSCs had higher immunofluorescent staining of the proliferation marker KI67 (Figure 5A, Supplementary Figure 5) and had higher sphere forming capacity (Figure 5B). To more directly establish a relationship between proliferation and reliance on FEN1, we treated highly proliferative FEN1i-sensitive GSCs with Paclitaxel (PTX), an inhibitor of mitosis and cell division, which stalled cell proliferation while maintaining viability. PTX treatment resulted in significant resistance to FEN1 inhibition (Figure 5C), indicating the importance of FEN1 to proliferative cells. To further show this relationship, we did the opposite and hyper-proliferated slow-dividing FEN1i-resistant GSCs using TP53 knockout, a common mutation found at GBM recurrence. Doing so sensitized the FEN1i-resistant GSCs to FEN1 inhibition (Figure 5D), further supporting the importance of FEN1 function in proliferative cells.

**Figure 5.**
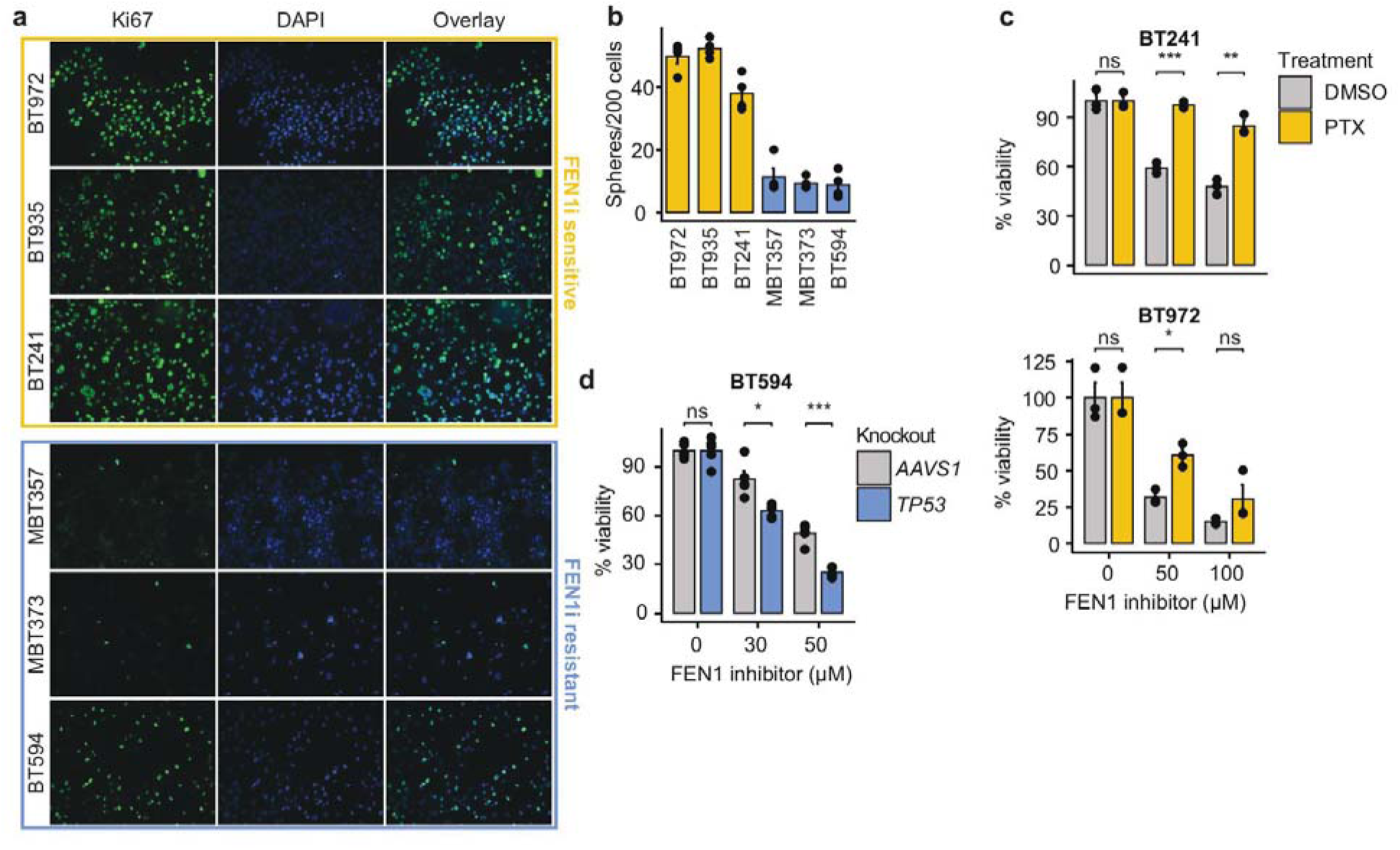
FEN1 dependency is associated with tumor cell stemness and proliferation. (A) Immunofluorescence staining for Ki67 and DAPI in FEN1i-sensitive (BT972, BT935, BT241) and resistant (MBT357, MBT373, BT594) GSCs. (B) Secondary sphere formation assay in FEN1i-sensitive and resistant GSCs. Data are represented as mean ± s.e.; n = 4 experimental replicates. (C) Cell viability assays of FEN1i-sensitive GSC lines cultured with or without PTX (cell cycle stalling agent) for 6 days, exposed to increasing concentrations of FEN1i. Data are represented as mean ± s.e.; n = 3 experimental replicates. (D) Cell viability assay of FEN1i-resistant GSCs with TP53 or AAVS1 (control) knockout, exposed to increasing concentrations of FEN1i. Data are represented as mean ± s.e.; n = 5 experimental replicates.

To more broadly characterize the role of FEN1 in GSC survival, we analyzed single-cell RNA sequencing (scRNA-seq) data from patient-derived GBM cell lines from the Abdelfattah et al. (2022)^15^ and Wang et al. (2022)^16^ datasets. At the gene level, FEN1 expression correlated strongly with key regulators of cell cycle progression, stemness, and DNA damage repair (Figure 6A, Supplementary Table 4). Consistent with our prior immunofluorescence findings, one of the strongest co-dependencies was MKI67, the gold-standard proliferation marker in GBM. Other proliferation-associated genes included PCNA, a master regulator of DNA replication, and CCNE2, which also promotes rapid cell cycle progression. Additionally, multiple genes involved in mitosis, chromosomal segregation, and spindle assembly—including TPX2, SPC25, DSN1, DTL, and NCAPG—were highly co-expressed with FEN1. From a stemness perspective, FEN1 correlated with key GBM stemness-associated genes, including EZH2, ATAD2, and HELLS, which epigenetically regulate cancer cell plasticity and self-renewal. Additionally, ASF1B and PBK, which contribute to stem-like properties and tumor progression, were highly co-expressed. FEN1 also showed strong associations with genes involved in DNA damage repair pathways, particularly homologous recombination, including BRCA1, BRCA2, RAD51AP1, and XRCC2. Genes within the Fanconi anemia (FA) pathway, such as BRIP1, FANCI, UBE2T, and USP1, were also highly co-expressed, further supporting a role for FEN1 in maintaining genomic stability. Finally, CHEK1, a master checkpoint kinase essential for DNA damage response, was among the top co-expressed genes.

**Figure 6.**
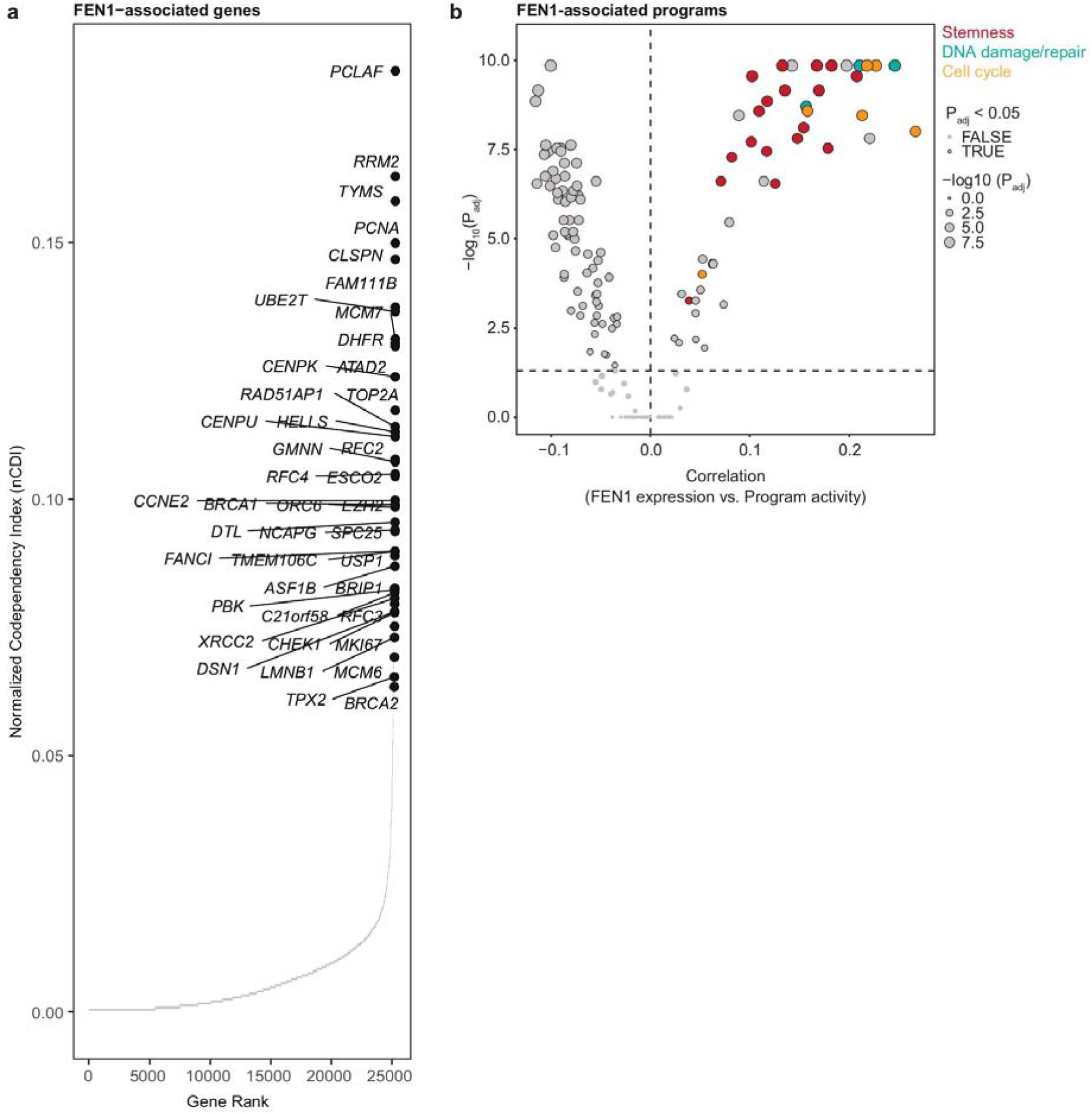
Single-cell analysis reveals that FEN1 expression correlates with key genes and gene programs involved in GBM proliferation, stemness, and DNA damage repair. (A) Co-dependency analysis of FEN1 expression, with co-expression represented by normalized co-dependency index. Key genes are highlighted and labeled, indicating strong co-expression with FEN1. (B) Gene program analysis showing correlation between FEN1 expression and program activity. Dot size corresponds to −log10 (adjusted P value), with larger dots indicating more statistically significant results. Dots are color-coded according to gene program category, and key programs are highlighted.

At the gene program level, FEN1 expression was again significantly enriched in programs related to cell cycle/proliferation, stemness, and DNA damage repair, further reinforcing its role in aggressive GBM biology (Figure 6B, Supplementary Table 5). Notably, proliferation and cell cycle programs, including MP1_CellCycle_G2M,^19^ MP2_CellCycle_G1S,^19^ MP3_CellCycle_HMG,^19^ CellCycle_Cancer,^22^ and HALLMARK_G2M_CHECKPOINT^49^ were highly correlated with FEN1 expression. Stemness programs, such as StemSig_Zhang2022,^24^ Nanog_Benporath,^26^ Sox2_Benporath,^26^ Human_Epithelial_ASC_Smith,^50^ and hESC_Bhattacharya^51^ were also enriched, highlighting FEN1’s association with GSC identity. Finally, DNA damage repair pathways, including HALLMARK_DNA_REPAIR,^49^ DNArepair_CancerSEA,^22^ and DNAdamage_CancerSEA^22^ were highly expressed in GBM cells expressing FEN1, consistent with its role in genomic maintenance.

These findings suggest that FEN1 is a central regulator of proliferation, stemness, and DNA damage repair in GBM, making it a promising therapeutic target, particularly in combination with DNA-damaging agents such as temozolomide.

## Discussion

Despite decades of progress in cancer biology and therapeutics, glioblastoma (GBM) remains one of the most treatment-refractory and deadly cancers, underscoring the urgent need for innovative therapeutic strategies. Functional genomics has begun to shed light on therapy resistance in GBM. MacLeod et al. previously implicated Fanconi anemia and homologous recombination pathways (including genes such as FANCA, C19orf40, MCM8, and MCM9) as contributors to temozolomide (TMZ) resistance in primary glioblastoma stem cells (GSCs).^10^ Similarly, Rocha et al. highlighted resistance pathways involving Sonic Hedgehog signaling, circadian rhythm genes, the NRF2 antioxidant response, and Wnt/β-catenin signaling, identifying genes such as CLCA2, PTCH2, FZD6, and CTNNB1 which promote cell survival when overexpressed, suggesting they could be targeted to improve TMZ efficacy.^52^ Despite these advances, the functional genetic regulators underpinning GSC resistance remain poorly understood, and actionable therapeutic targets are scarce. In this study, we conducted the first functional genetic screening in patient-derived GBM models to investigate essential gene modulators of standard of care (SoC), uncovering flap endonuclease 1 (FEN1) as a critical driver of survival and TMZ resistance in an aggressive subset of GBMs. Our findings position FEN1 as a cancer-selective therapeutic target with potential to enhance the efficacy of SoC in aggressive, treatment-refractory GBM.

Our chemoradiogenetic screening revealed that glioblastoma stem cells (GSCs) display diverse and condition-specific dependency profiles, underscoring the complexity of therapy resistance mechanisms. We propose that resistance may arise from broad, essential genetic programs responsible for tumorigenesis, growth and survival which have thus far eluded genomic efforts. The preferential sensitization of FEN1i-sensitive GSCs to TMZ by FEN1 inhibition supports a model in which genes essential for tumorigenic properties further shield tumor cells from genotoxic therapies. This finding highlights the potential of targeting fundamental tumorigenic dependencies to overcome resistance and should prompt further exploration into the involvement of essential genes in these processes.

Targeting cancer stem cell properties in GBM has long been challenging due to the homology with healthy neural stem cells (NSCs). Additionally, clinical outcomes of NSC toxicity are uncertain, and while NSCs remain largely quiescent post-development, ample evidence supports their roles in aging, plasticity, disease, and regeneration of the nervous system. Despite these challenges, several preclinical and clinical strategies have aimed to target stem-like properties in GBM, including BMI1 and EZH2 inhibitors and CD133 CAR T cells.^53,54^ Our study contributes to this effort by advancing the understanding of GSC-specific dependencies, particularly those related to DNA damage response pathways. These pathways, which maintain mutational tolerance and genomic stability, are critical for maintaining plasticity and stemness of cancer stem cells. Reliance on these proteins presents a targetable dependency enriched by exposure to genotoxic therapies, and FEN1 emerges as a key dependency within this framework, representing a novel and actionable vulnerability in GSCs.

In addition to deciphering cancer-specific stem cell biology, the field is further complicated by the temporal evolution GBM undergoes throughout the course of therapy. It has been well established that recurrent GBM is markedly different from the primary disease, being more aggressive and more resistant to a broad range of therapeutics, and a lack of scientific discovery in recurrent GBM models likely contributes to the failures seen across clinical trials.^55,56^ Thus, therapeutic strategies that maintain or enhance efficacy in the highly proliferative, therapy resistant phenotype characteristic of recurrent GBM are crucial. Additionally, upon acquiring this resistant phenotype, patients with recurrent GBM often refuse secondary TMZ treatment due to a lack of efficacy. Identifying therapeutic strategies that are effective against recurrent GBM and further address this resistance to therapy may improve the efficacy of secondary TMZ and allow it to further extend patient survival.

While our study provides compelling evidence for FEN1 as a novel therapeutic vulnerability in GSCs, we are limited by a lack of in vivo validation, which remains a critical next step in translating these findings toward clinical impact. Notably, the lack of blood-brain barrier (BBB)-penetrant FEN1 inhibitors poses a current limitation to such studies; however, this challenge also represents an exciting and necessary opportunity for future drug development. Given the urgent clinical need for novel, effective treatments in GBM—particularly for recurrent, treatment-resistant disease—the development of BBB-penetrant FEN1-inhibitors could enable preclinical in vivo studies and ultimately catalyze the translation of our findings into viable clinical strategies. As such, FEN1 represents both a promising therapeutic target and a foundation for future innovations in GBM drug discovery.

In summary, our study highlights FEN1 as a pivotal driver of GSC survival and therapy resistance, with dual roles in maintaining tumorigenic properties and mediating genotoxic therapy resistance. By targeting this dependency, we provide a foundation for novel therapeutic strategies aimed at disrupting critical vulnerabilities of stemness in GBM, particularly in its most aggressive and treatment-resistant forms.

## Acknowledgements

We thank all members of the laboratories led by Drs. Sheila K. Singh, Sachin Katyal, Jason Moffat and Deena Gendoo. S.K.S holds a Senior Canadian Research Chair in Human Cancer Stem Cell Biology and J.M holds a Canada Research Chair in Functional Genomics of Cancer.

## Author contributions

Conceptualization: B.A.B., C.R.C., C.V., J.M., S.K. and S.K.S.; Resources: J.M., S.K. and S.K.S.; Methodology, investigation and validation: B.A.B., C.R.C., D.M., A.P., V.M.S., M.Singh, A.S., A.T., N.M., M.B., A.A., P.M., K.R.B., D.T., W.M., S.S., Y.S., M.Subapanditha, D.M.A.G., C.V.; Software and formal analysis: B.A.B., C.R.C., N.M., M.B., K.R.B., D.M.A.G.; Visualization: B.A.B. and C.R.C.; Writing: B.A.B. and C.R.C., with input from other authors; Project administration and supervision: C.R.C., S.K.S., S.K., J.M. and D.M.A.G.; Funding acquisition: S.K.S., S.K. and J.M. All authors read and approved the manuscript.

## Supplementary Data

**Supplementary Table 1.** Antibodies used for immunoblotting.

**Supplementary Table 2.** Bayes factors for CRISPR-Cas9 dropout screens, related to figure 1B.

**Supplementary Table 3.** Normalized gRNA read counts for chemoradiogenetic CRISPR-Cas9 screen in GBM model BT935, related to figure 2B.

**Supplementary Table 4.** Genes associated with FEN1 expression in scRNA-seq analysis, related to figure 6A.

**Supplementary Table 5.** Programs associated with FEN1 expression in scRNA-seq analysis, related to figure 6B.

**Supplementary Figure 1.**
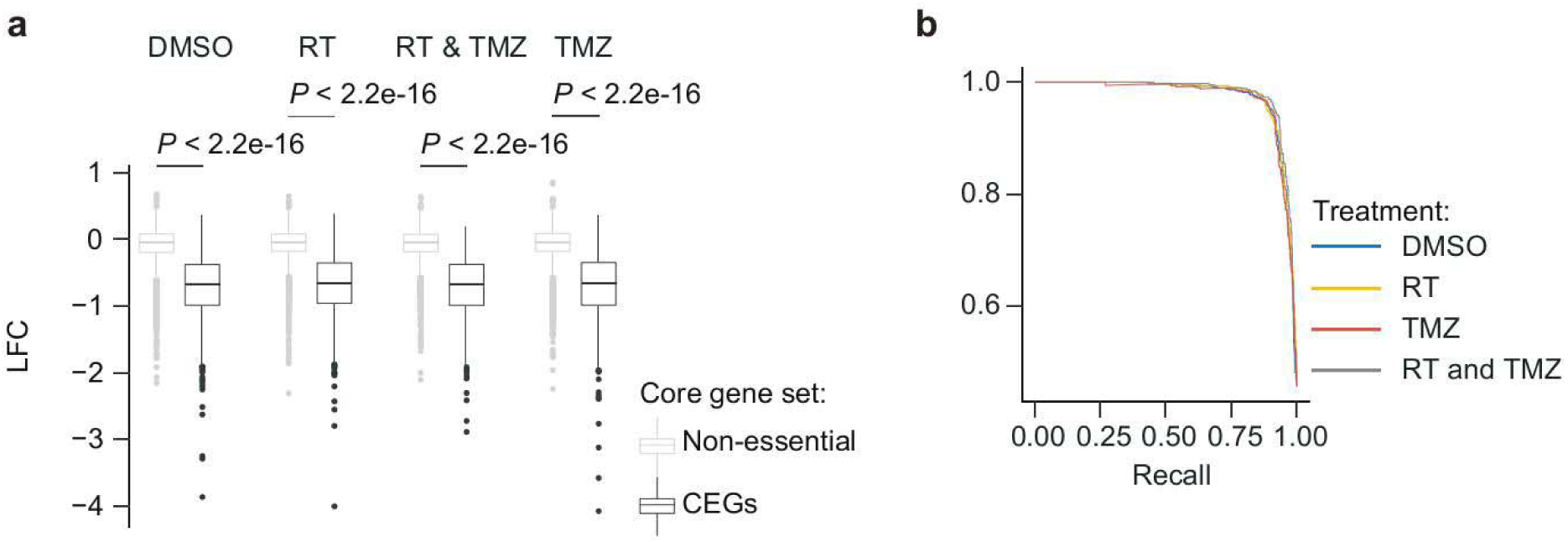
Quality control analysis of CRISPR-Cas9 dropout and chemoradiogenetic screens in GBM model BT935, related to figure 1B and 2B. A) Gene-level LFC of CEGs and non-essential genes across treatment arms. All data presented from n = 3 biologically independent replicates. P values are from unpaired Student’s t tests. B) Precision-recall plots indicate high performance of screens across treatment arms. C) Number of essential genes (FDR < 0.05) identified in screens across treatment arms. D) Number of conditional genetic interactions identified in screens across treatment arms (FDR < 0.05).

**Supplementary Figure 2.**
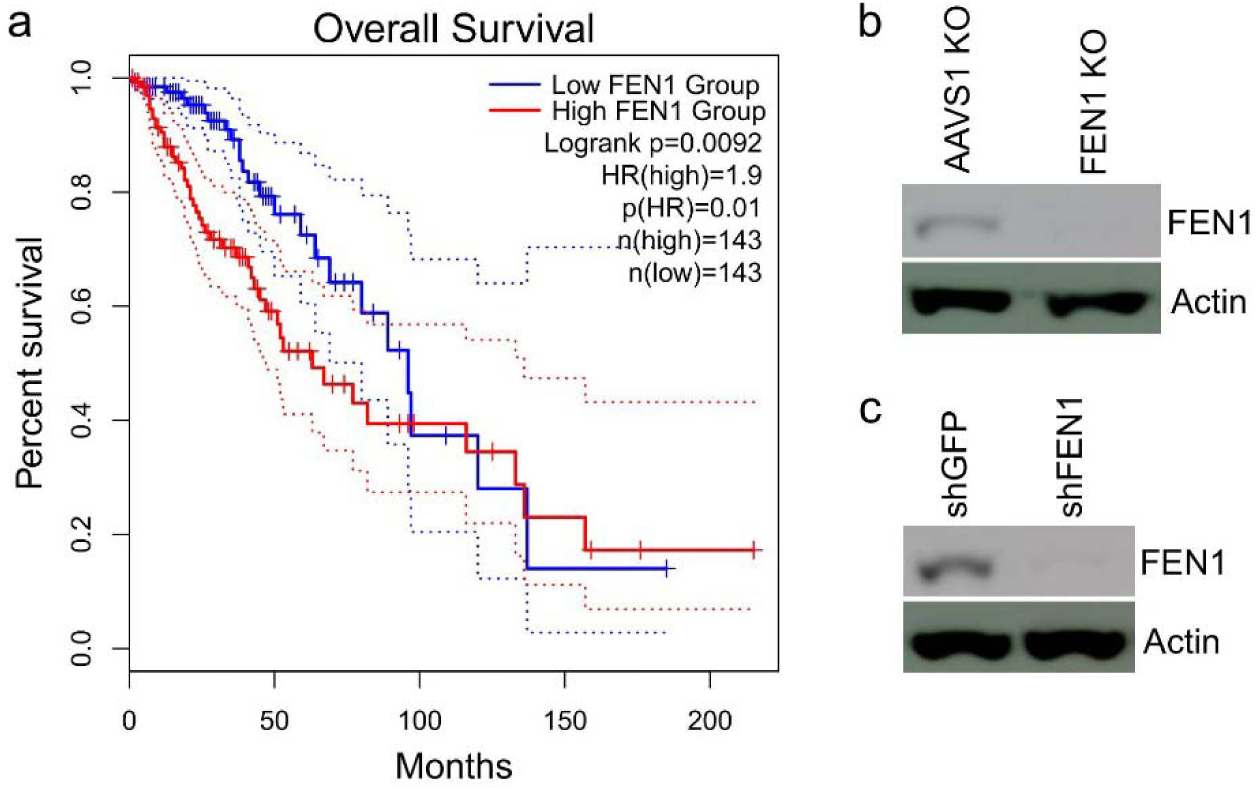
A) Kaplan–Meier survival analysis of GBM patients with high- and low-FEN1 expression, from TCGA data. B) Immunoblotting of FEN1 protein in GSCs after knockout of FEN1 or AAVS1 (control), related to figure 2C and 2D. C) Immunoblotting of FEN1 protein in GSCs after knockdown of FEN1 or GFP (control), related to figure 2E.

**Supplementary Figure 3.**
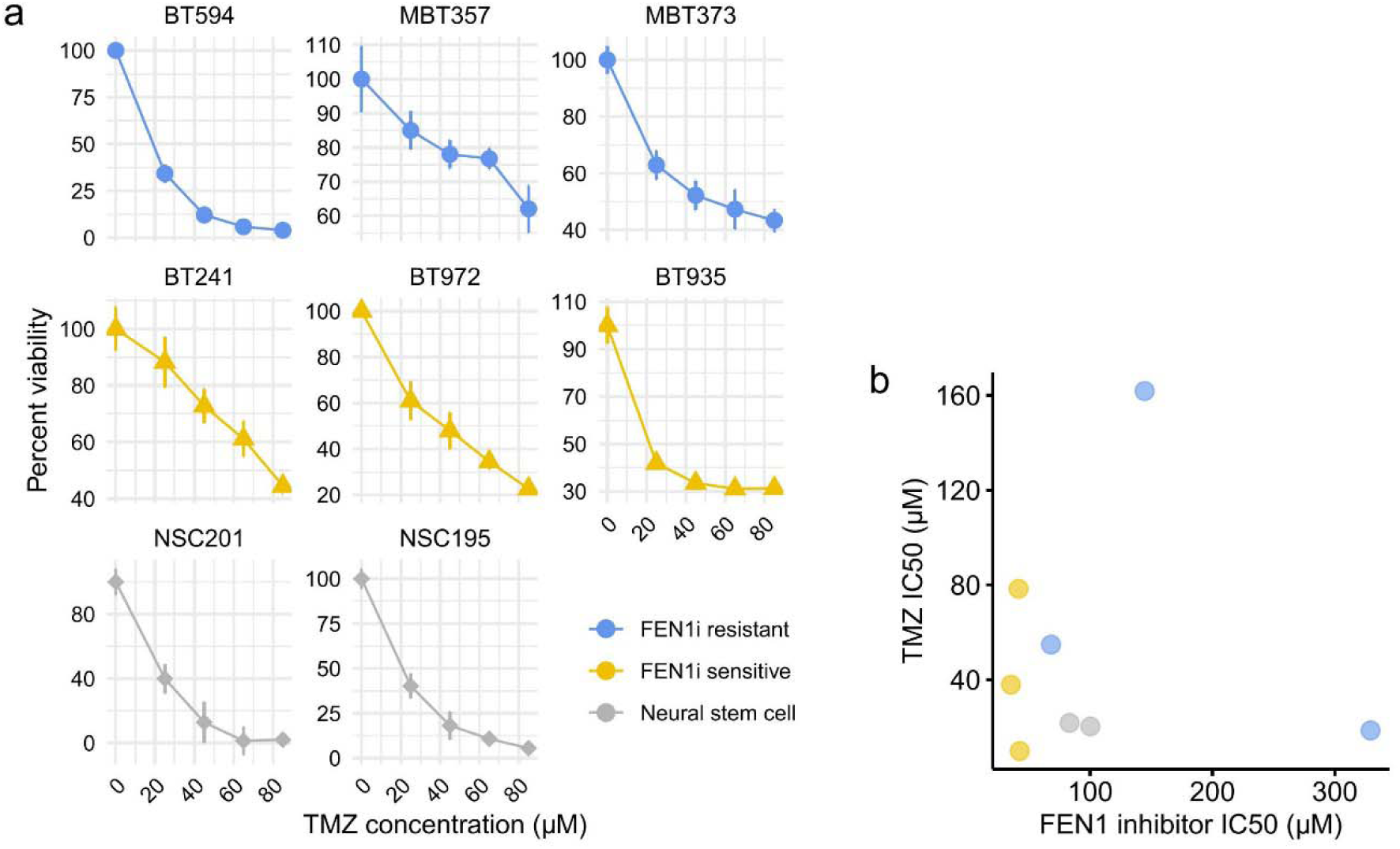
A) Cell viability assays of GSC lines exposed to increasing concentrations of TMZ inhibitor over 7 days, related to figure 3. Data are represented as mean ± s.e.; n = 4 experimental replicates. B) Plot of TMZ and FEN1i IC50 values, related to figure 3.

**Supplementary Figure 4.**
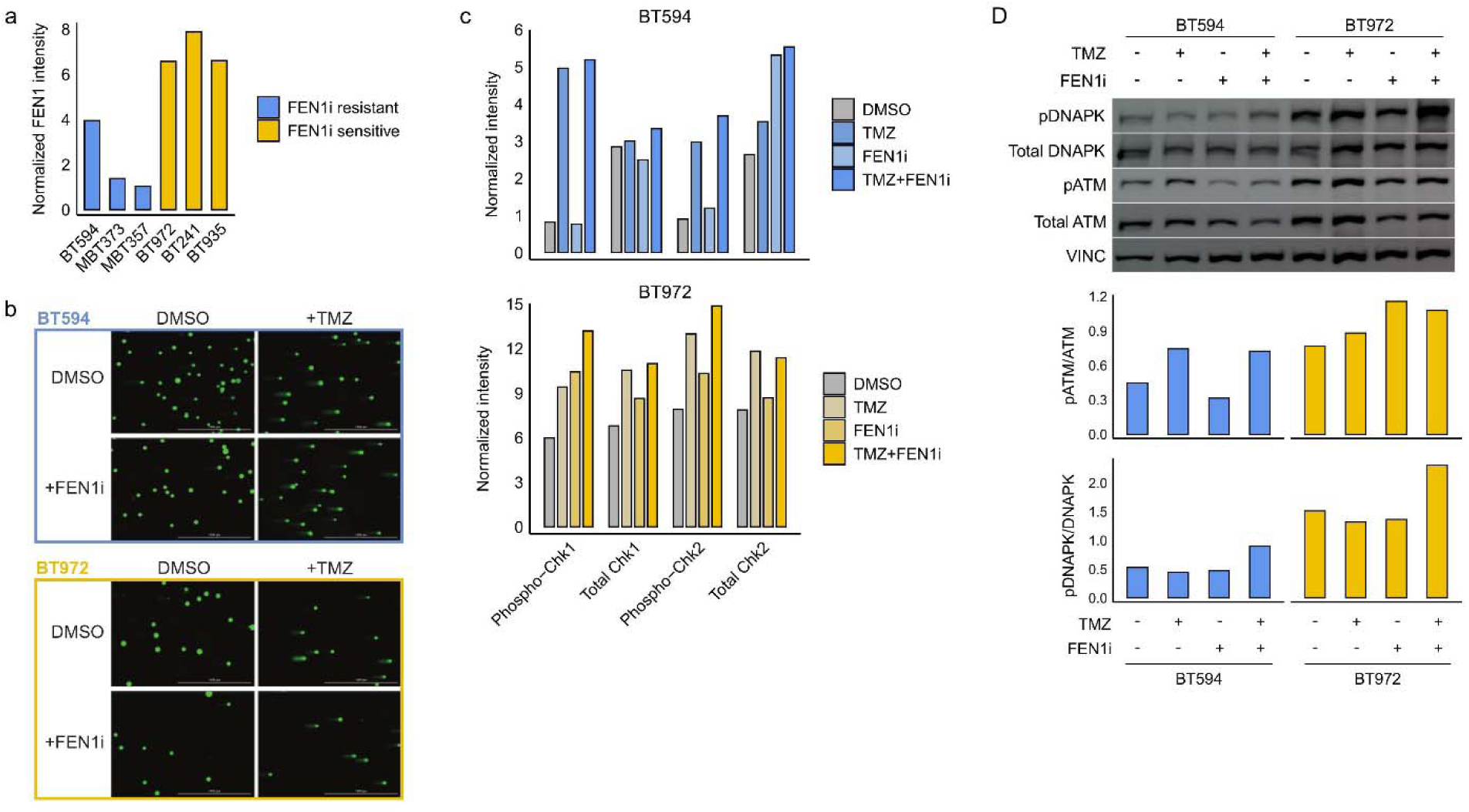
A) Normalized intensities of FEN1 protein from immunoblotting related to figure 4A. B) Representative images of comet assays of FEN1i-resistant (BT594) and sensitive (BT972) GSCs exposed to DMSO alone (control), TMZ (200 µM), FEN1i (30 µM) or combination (TMZ/FEN1i, 200 µM/ 30 µM). C) Normalized intensities of phospho-Chek1, total Chek1, phospho-Chek2 and total Chek2 protein from immunoblotting related to figure 4E and 4F. D) Immunoblotting for phospho-ATM, total ATM, phospho-DNAPK and total DNAPK protein normalized to vinculin control.

**Supplementary Figure 5.**
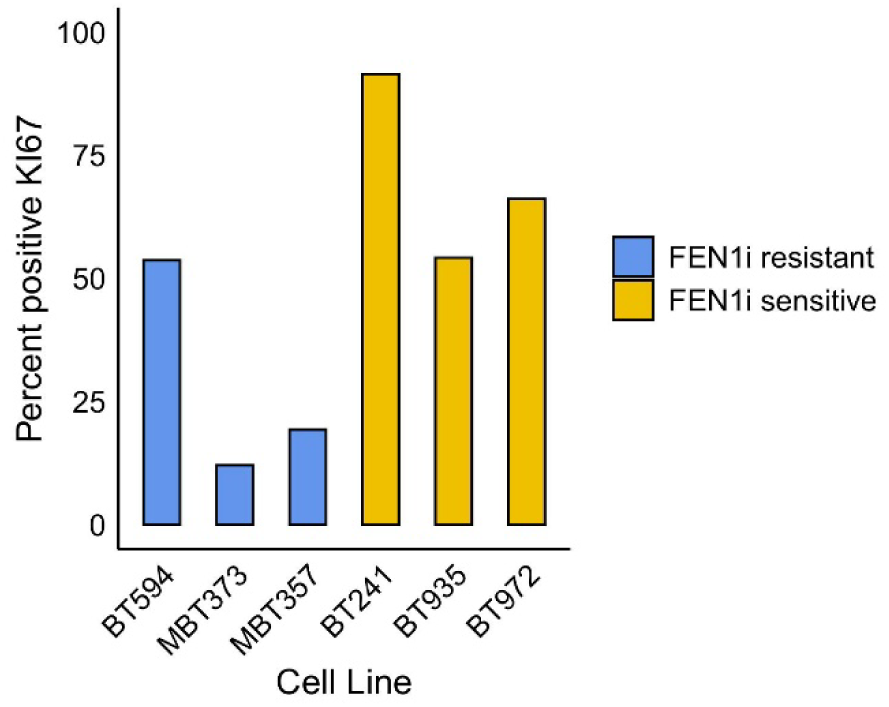
Quantification of KI67 staining, as percent-positive for KI67, related to figure 5A.

